# Single cell network analysis with a mixture of Nested Effects Models

**DOI:** 10.1101/258202

**Authors:** Martin Pirkl, Niko Beerenwinkel

## Abstract

**Motivation:** New technologies allow for the elaborate measurement of different traits of single cells. These data promise to elucidate intra-cellular networks in unprecedented detail and further help to improve treatment of diseases like cancer. However, cell populations can be very heterogeneous.

**Results:** We developed a mixture of Nested Effects Models (M&NEM) for single-cell data to simultaneously identify different cellular sub-populations and their corresponding causal networks to explain the heterogeneity in a cell population. For inference, we assign each cell to a network with a certain probability and iteratively update the optimal networks and cell probabilities in an Expectation Maximization scheme. We validate our method in the controlled setting of a simulation study and apply it to three data sets of pooled CRISPR screens generated previously by two novel experimental techniques, namely Crop-Seq and Perturb-Seq.

**Availability:** The mixture Nested Effects Model (M&NEM) is available as the R-package mnem at https://github.com/cbgethz/mnem/.

**Supplementary information:** Supplementary data are available.online.

## 1 Introduction

Understanding heterogeneous diseases like cancer on a molecular level is challenging, but also crucial for the improvement and development of therapies. Molecular intra-tumor heterogeneity is an important factor for cancer treatment (Sun, 2015, Prasetyanti and Medema, 2017). Treatments often assume cancer to be homogeneous across cells. However, if different cell types are resistant to different treatments, the success of current treatment strategies is limited.

A key component of the molecular landscape are signaling pathways and how they are causally wired in healthy and diseased cells. De-regulation of pathways in diseased cells is prevalent (Mao, 2012, Giancotti, 2014) and to study this de-regulation, different mathematical methods have been developed. Several different algorithms have been proposed to analyze causal interactions of genes from different types of data (Friedman *et al.*, 2000; Nachman *et al.*, 2004; Margolin *et al.*, 2006; Kalisch and Bühlmann, 2007). Nested Effects Models (NEM, Markowetz *et al.*, 2005, 2007) infer pathways from perturbation data. In each experiment, one protein in the pathway is knocked down and a multi-trait read-out is produced, e.g., gene expression or cell imaging data (Siebourg-Polster *et al.*, 2015). If the expression of one gene changes during the knock-down compared to the unperturbed control, the knock-down has an effect on the gene and the gene responds to the knockdown. If the genes responding to the knock-down of protein B are a subset of the genes responding to the knock-down of protein A, NEMs will place A upstream of B in the pathway and a causal edge A to B is inferred.

NEMs have been successfully applied to different biological data sets to infer the causal network of signaling pathways (Markowetz *et al.*, 2005; Froehlich *et al.*, 2009; MacNeil *et al.*, 2015). Several extensions of NEMs have been developed, e.g. to account for hidden variables (Sadeh *et al.*, 2013). Epistatic Nested Effects Models (Pirkl *et al.*, 2017) systematically infer epistasis from double knock-down screens. Boolean Nested Effects Models (Pirkl *et al.*, 2016) make use of arbitrary combinations of knock-downs and knock-ins per experiment to infer a full boolean network and additionally integrate literature knowledge. Dynamic Nested Effects Models (Anchang *et al.*, 2009; Froehlich *et al.*, 2011) infer the rate of the signal flow within the network from time series data, while Hidden Markov Nested Effects Models (Wang *et al.*, 2014) model the evolution of the network itself during a time course. NEMix (Siebourg-Polster *et al.*, 2015) introduces a hidden variable to account for unobserved pathway activation.

The arrival of single-cell technologies provides new opportunities to improve resolution and account for heterogeneity in a population of cells. Pooled CRISPR screens enable gene expression measurements for thousands of cells with each cell having been the target of a CRISPR modification, i.e. a knock-down (Dixit *et al.*, 2016; Datlinger *et al.*, 2017). However, the heterogeneity in cell populations measured with single-cell technologies remains an open problem and there is a need for methods tailored to this new type of data.

Motivated by evidence, that causal signaling pathways can be differently wired in sub-populations of cells (Gaudet and Miller-Jensen, 2016), we introduce a mixture model, which simultaneously infers different sub-populations of cells across knock-downs and a causal network of the perturbed genes (Fig. 1). Cells are not hard clustered, but soft, such that each cell has a certain probability of being generated by each network (component). This probability defines how much a cell contributes to the network inference for each component.

**Figure 1:**
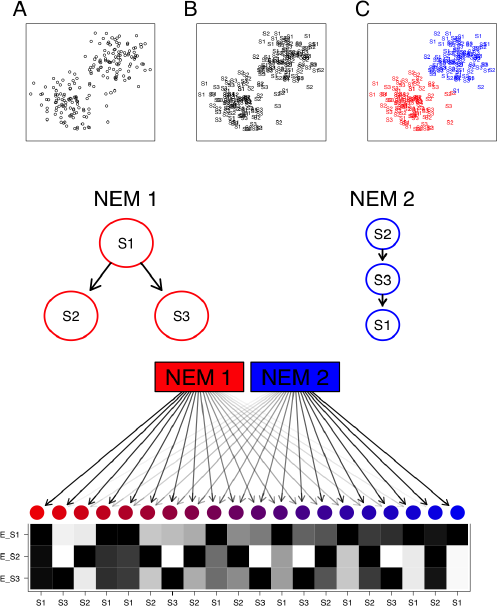
A schematic example of an M&NEM. (Top) Two dimensional projection of single cells (A), labeled for known knock-outs of signalling genes S1, S2 and S3 (B) and colored for their two clusters (C) on which the two different causal networks are based. (Middle) Comparing two different networks (NEMs) according to the clustering. S1, S2 and S3 are the perturbed genes and the edges denote how they causally influence each other. (Bottom) 20 noisy cells attached to each NEM according to their responsibilities with their respective data patterns (black for effect and white for no effect). Each row shows the effect pattern for the respective S-gene, e.g. E_S1 shows the effects of S-gene S1 for different cells. The colors and arrow transparencies depict the strength of attachment to the respective NEM. For example the bright red cell to the far left is attached to NEM 1 with responsibility 100%. Thus the pattern for the cell is the expected data pattern according to NEM 1 without any noise. The cells in the center are a mix of expected data patterns of cells for NEM 1 and NEM 2.

We show that Mixture Nested Effects Models (M&NEMs) work well in the controlled setting of a simulation study and apply our method to three data sets from two different pooled CRISPR screens based on Crop-Seq (Datlinger *et al.*, 2017) and Perturb-Seq (Dixit *et al.*, 2016). In those screens thousands of cells were pooled and each transfected with a different sgRNA to knock-out a specific gene. Gene expression data was generated by single cell RNA-Seq. For the Crop-Seq screen we concentrated on one data set investigating the T-cell receptor pathway in the T-Cell leukemia derived Jurkat cell line and key regulators DOK2, EGR3, LAT, LCK, PTPN6, PTPN11 and ZAP70. From the Perturb-Seq screen we model the causal interplay of cell cycle genes in one data set and transcription factors in another data set. Both data sets of the Perturb-Seq screens are derived from K562 leukemia cells.

## 2 Overview

In this section we review the original Nested Effects Model and extend it to a mixture of NEMs. Furthermore we discuss identifiability and propose a method for model selection to prevent over fitting.

### 2.1 Nested Effects Model

A Nested Effects Model (NEM) is parametrized by an adjacency matrix Φ ∊ *M_n×n_*({0, 1}) for the directed acyclic graph (DAG) representation of the signaling graph with perturbed genes as nodes (S-genes) and an adjacency matrix Θ ∊ *M_n×m_*({0, 1}) for the attachments of the different features from the data (E-genes), e.g., genes from gene expression data. *θ_ij_* = 1, if E-gene j is attached to S-gene *i*. Each column of Θ has at most one non-zero entry, because NEMs make the assumption that each E-gene can have at most one parent. Similar to Tresch and Markowetz (2008) we add a null S-gene, which predicts no effects to account for uninformative features.

We calculate the expected E-gene profiles for a given model (Φ, Θ) as the matrix product

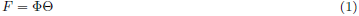

with *f_ij_* the predicted state of E-gene *i* in knock-down *j*.

Let 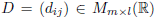 be the raw data matrix of the perturbation experiments and 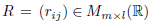 the log ratio matrix with perturbed cells indexing the columns and observed genes indexing the rows,

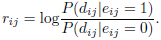

with *e_ij_* the unknown state of E-gene *i* in knock-down *j*. As in Tresch and Markowetz (2008) we can write the log likelihood ratio of a given model (Φ, Θ) and the null model *N*, which predicts no effects, as

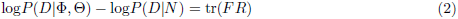

where tr denotes the trace of a quadratic matrix. However, *F R* is only quadratic if the data includes only one cell per knock-down, i.e. *l* = *n*. Hence, the data has to be summarized beforehand, e.g., by taking the average over all experiments with the same knock-down (replicates).

### 2.2 Mixture Nested Effects Model

Instead of inferring a single network Φ and E-gene attachments Θ from the whole data set as in the previous section, we formulate a mixture, which infers several networks with unique attachments and different sub-populations of cells.

The model parameters for a mixture of *K* components (Φ, Θ) are

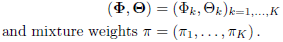

Given a component (Φ*_k_*, Θ*_k_*) we calculate the expected knock-down profiles for all single perturbations using Eq. 1 as

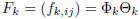

with *f_k,ij_* the expected value of E-gene *j* under the perturbation of S-gene *i* in component *k*.

The log ratio profile of all cells given component *k* is

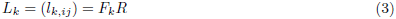

and the log likelihood ratio of component *k* is

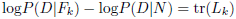

Let *Z* ∊ *M_K×l_*({0, 1}) be a matrix for the hidden cell attachments to the components. *z_ki_* = 1, if cell *i* belongs to component *k*. Each column of *Z* has exactly one non-zero entry. The distribution of *Z* is defined by the mixing coefficients *π_k_* as

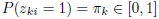

for all *i* ∊ {1, …, *l*} with *π* = (*π_1_, …, π_K_*) and 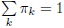.

**Log likelihood of the mixture**. For model optimization we choose a maximum likelihood (ML) approach using the log likelihood ratios similarly to the formulation for a single mixture component,

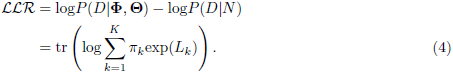

The full derivation of the likelihood ratio is in Eq. S1 of the supplement.

### 2.3 Inference with a Expectation maximization algorithm

We developed an Expected Maximization scheme (Dempster *et al.*, 1977) for inference.

**E step**. Let *π*, (**Φ, Θ**) be the current parametrization of our mixture model. We calculate *L_k_* from Eq. 3 with

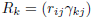

with cell and component specific weights *γ_kj_* substituted for *R* for every component *k* and subsequently the responsibilities (supplement, Eq. (S2))

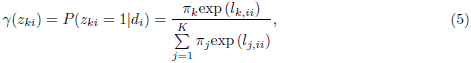

which we summarise in

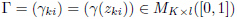

and the log likelihood ratio (Eq. 4).

***M*_Φ_ step**. We update *π* with

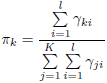

**Φ** remains fixed and we estimate **Θ** by their maximum a posteriori attachment to each S-gene. For this we use the known perturbation map *ρ* = (*ϱ_ij_*) with *ϱ_ij_* = 1, if cell j has been perturbed by a knock-down of S-gene i. We compute the fit of every E-gene to every S-gene

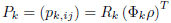

and set

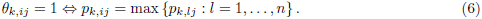

We alternate between the *E* step and the *M*_Θ_ step until the log likelihood ratio in Eq. 4 converges.

**M step**. Given Γ, we optimize each component (Φ*_k_*, Θ*_k_*) with respect to *R_k_*. We maximize the log likelihood ratio defined in Eq. 2 to find new optimum 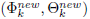 in the following way.

We optimize each individual component with a natural extension of the module network approach by Froehlich *et al.* (2008). We cluster knock-downs, averaged over cells, into groups of size n (e.g. *n* = 5) and perform a local neighborhood search on each group. In the local neighborhood search we evaluate each edge for absence and presence and check whether a change in status improves the log likelihood ratio and change the edge which improves it most. We combine the inferred sub-networks to one large network including all S-genes and use it as the initial network for a local neighborhood search on the full set of S-genes. During the optimization of *Φ_k_*, we estimate *Θ_k_* as in Eq. 6 before we calculate the log likelihood ratio.

We alternate between the *E*, *M*_Θ_ and *M* steps until the the log likelihood ratio in Eq. 4 converges. To increase the probability of convergence to a global optimum, the EM algorithm is initialized several times with random responsibilities between 0 and 1.

### 2.4 Model identifiability

In the case of the original NEMs, two NEMs Φ_1_ and Φ_2_ are identical if and only if they have equal transitive closures, i.e. they produce identical data. This identity still holds for each component of a mixture of NEMs. However, mixture NEMs have additional identifiability issues.

In general, two M&NEMs are not distinguishable, if they generate the same data. Let *F = (F_1_, …, F_m_*) be the expected data pattern for M&NEM A and 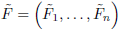 the expected data pattern for M&NEM B. If each column *f_v_* of *F* is included in 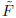 and each column 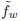 of 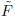 is included in *F*, A and B are not distinguishable.

Fig. 2 shows a schematic example for two identical mixtures (A,B) with different components. For convenience of this example we assume an a posteriori hard clustering of the cells to the components and equal attachments Θ_1_ = Θ_2_. For two cell clusters we compute an optimal mixture of two NEMs (A). However, if we divide the same data into two different clusters, we compute an optimal mixture of NEMs (B), which differs from A. Nevertheless, both mixtures perfectly explain the same data and are therefore indistinguishable from each other.

**Figure 2:**
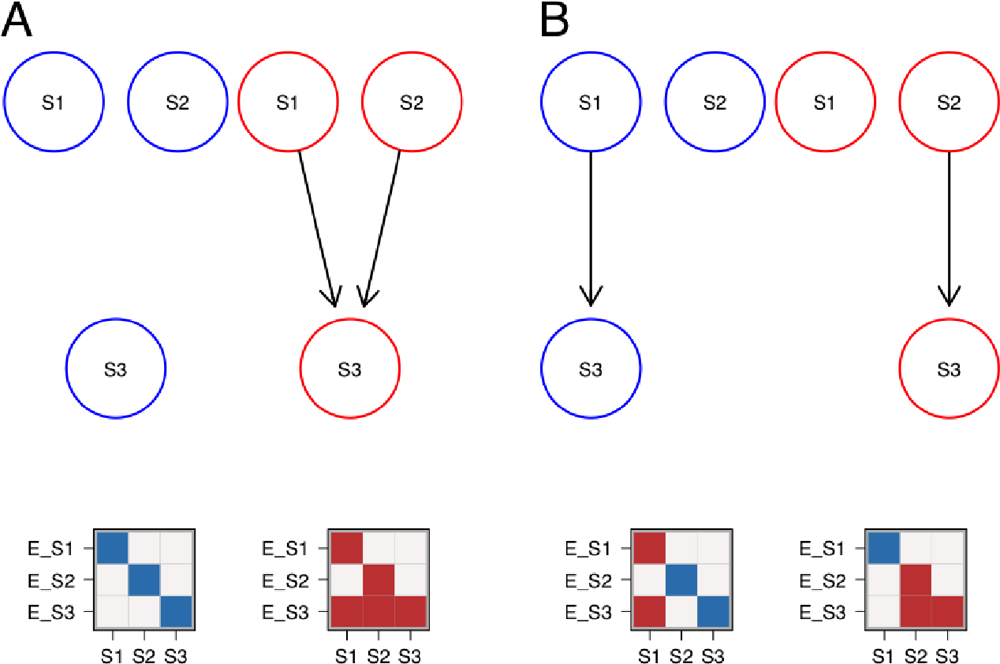
Example for non-identifiability of two M&NEMs. (**Top**) Mixture of two components (blue, red) with their respective attached cells (data) below. Dark areas are effects and light areas are no effects. Each column of the data is a cell and each row is the effect pattern for gene E_X attached to S-gene X. (**B**) Identical mixture to **A**. The first columns (cells) of each cluster have been switched. Overall the data stays the same, but the network components changed. However, they still yield a perfect fit to the data.

### 2.5 Model selection

In a typical situation for M&NEMs we do not know the correct number of components *K*. To prevent over fitting and enforce sparsity to the solution, we choose the optimal *K* via a penalized log likelihood ratio, penalizing complex and redundant network structures in a similar fashion as Froehlich *et al.* (2007). For each *K* ∊ {1, …, 5} we infer an optimal solution using the EM. Then we score each of the five solutions with a penalized log likelihood ratio, which we define as

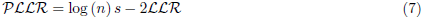

with a complexity parameter *s*, model log likelihood ratio *LLR* (Eq. 4) and the sample size *n* (number of cells). We define *s* for a mixture of *K* components as

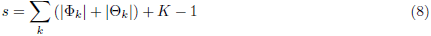

with number of edges of an adjacency matrix *A* denoted by *|A|*. Thus the number of parameters *s* are all edges in the graphs of *_k_* and *_k_* plus one less than the number of mixture weights, since the last weight is determined by the others. Finally we choose the solution, which minimizes the penalized log likelihood ratio. Fig. 3 shows the raw and the penalized log likelihood ratio as functions of the number of components for the data sets in our application.

**Figure 3:**
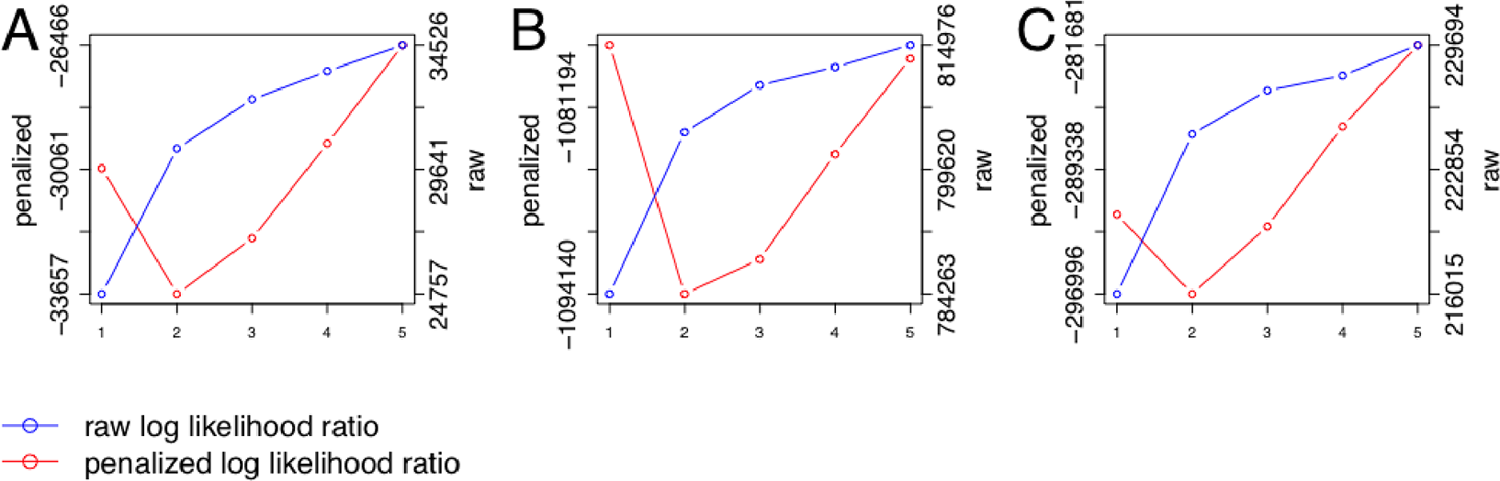
Penalized log likelihood ratio (red, left y-axis) in comparison with the raw log likelihood ratio (blue, right y-axis) as functions of number of components for the Crop-Seq regulators of the T-Cell receptor (A) and the Perturb-Seq cell cycle genes (B) and transcription factors (C).

### 2.6 Effect log-odds

We calculate log odds for the effects analogous to Siebourg-Polster *et al.* (2015). Let *d_ij_* be the normalized count value for gene *i* and cell *j*. Cell *j* was perturbed by a knock-down of gene *k*. We estimate the empirical distribution function *F*_0_ of the normalized control counts for gene *i* and the empirical distribution function *F_k_* of the normalized counts from cells perturbed by *k* for gene *i* and calculate the log odds by

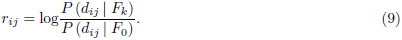

If the E-gene shows a clear effect in the cell, *r_ij_* will be greater than zero and if it shows no effect, it will be less than or equal to zero.

We remove E-genes with a standard deviation smaller than the global standard deviation over the whole data set, i.e. E-genes which have small log odds apart from outliers.

## 3 Simulations

We showed that M&NEMs work well in simulations under reasonable conditions. For *n* ∊ {3, 5, 10, 20} S-genes and *K* ∊ {1, 2, 3, 4, 5} we drew random mixture weights *π* and component(s) (**Φ, Θ**) as the ground truth. We simulated 1000 cells overall, two E-genes per S-gene and 10% uninformative E-genes. The simulated data were log odds with added Gaussian noise around −1 for no effect and 1 for effect. Fig. 4 shows the result of 100 runs and Gaussian noise *N* (0, *σ*) with *σ* ∊ {1, 2.5, 5}.

**Figure 4:**
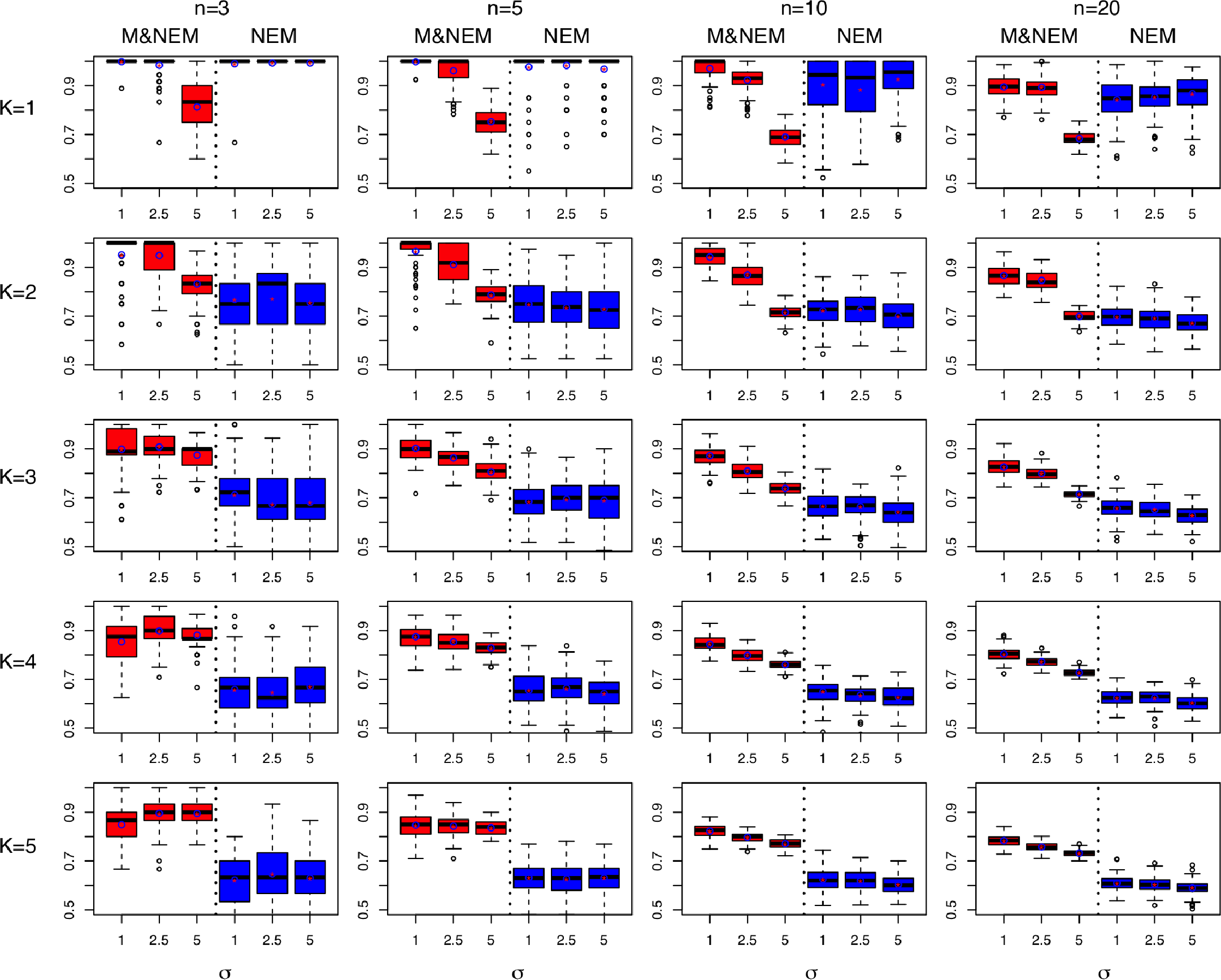
Comparison of M&NEMs and NEMs in a simulation study. The rows show results for components *K* ∊ {1, 2, 3, 4, 5}. The columns show results for number of S-genes *n* ∊ {3, 5, 10, 20}. Each box plot shows the accuracy of M&NEM (red) and NEM (blue) for three different noise levels *σ*. The y-axis is cutoff at 50% (=random). In addition to the median we also added the average (red star in blue circle).

We computed accuracy from similarity of the ground truth **Φ** = (Φ_1_, …, Φ*_K_*) and the inferred optimum 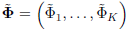. That is, we check how accurately we find a column from the ground truth **Φ** in the inferred optimum 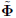 and vice versa with the following score,

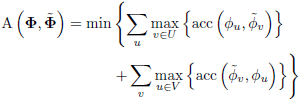

where *U* and *V* are the sets of columns with the same perturbation as *u* and respectively *v*, *ϕ_i_*, 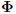 as the columns of **Φ** respectively 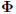 and

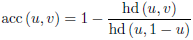

with the hamming distance hd.

The simulations show that M&NEMs can identify the ground truth with high accuracy for reasonable noise levels and is still robust in settings with high noise over a varying number of components and S-genes. The accuracy for *K* and the mixture weights are shown in Fig. S1-S2 of the supplement.

## 4 Application to pooled single cell CRISPR screens

In our application of M&NEM we analyze three data sets which combine pooled CRISPR screening with single cell RNA-seq readouts. One data set was generated with Crop-Seq (Datlinger *et al.*, 2017) and the other two with Perturb-Seq (Dixit *et al.*, 2016).

### 4.1 CRISPR droplet sequencing (Crop-Seq)

Datlinger *et al.*, 2017 combined pooled CRISPR screening with single-cell RNA sequencing to produce gene expression count data on the single-cell level. They showed the validity of their method with an analysis of T-cell receptor (TCR) activation in Jurkat cells. We downloaded the processed CROP-seq data from the NCBI GEO database (Edgar *et al.*, 2002, GSE92872). We reduced the data to stimulated cells and genes, which have a median count number of *>* 0 over the remaining cells. We normalized the count data to counts per 10000, i.e. we divided each count by the sum of counts of its respective column and multiplied by 10000. Next we took the log of the normalized counts plus a pseudo count of 0.5 and calculated log odds (Eq. 9).

As a set of knock-outs we concentrated on S-genes involved in T-Cell receptor activity as in Fig. 2, h of Datlinger *et al.* (2017), namely: DOK2, EGR3, LAT, LCK, PTPN6, PTPN11 and ZAP70. This leaves us with a population of 535 unique cells and 663 E-genes. Fig. 5 shows the result for the highest scoring result with *K* = 2. Around 58% of cells are assigned to the red network and 42% to the blue. M&NEM confirms key down-stream regulators LCK and LAT (Datlinger *et al.*, 2017, Fig. 2, h). However, ZAP70 is placed more upstream especially in the blue network. DOK2 on the other hand is correctly placed as an upstream regulator in the red network (Datlinger *et al.*, 2017, Fig. 2, h), but placed right downstream of everything else in the blue one, hinting at an altered causal role of DOK2 in the smaller cell population. PTPN6 and PTPN11 are placed as the main regulators in the red respectively blue network.

**Figure 5:**
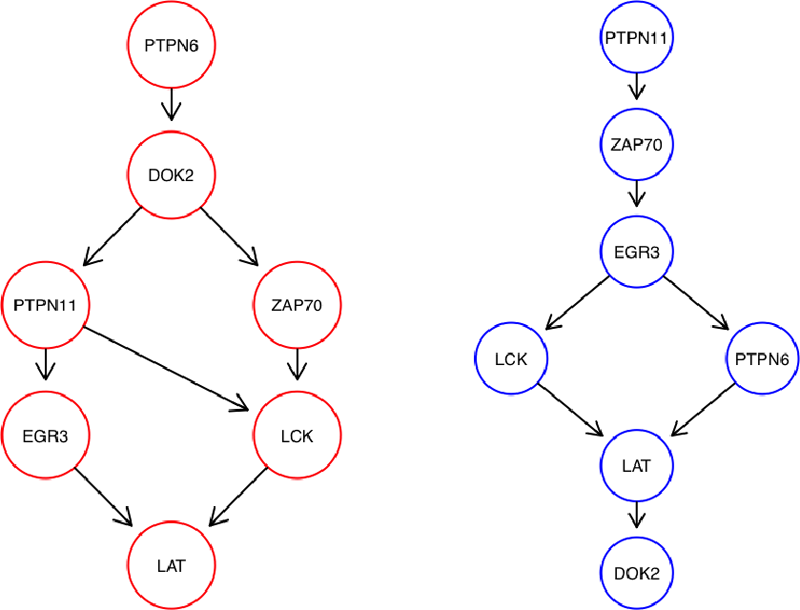
Optimal mixture found for the Crop-Seq data set (*K* = 2) with mixture weights 58% (red) and 42% (blue).

A posteriori a majority of 310 cells are attached to the red network. However, for DOK2 and PTPN6 the majority of cells for each knock-out are attached to the blue network, which explains the relatively high mixture weight of 42%, The responsibilities for each network are almost binary, 100% respectively 0% (Fig. 6, A). This is almost equivalent to a hard clustering of the cells, i.e. there is virtually no uncertainty of the cell attachments.

**Figure 6:**
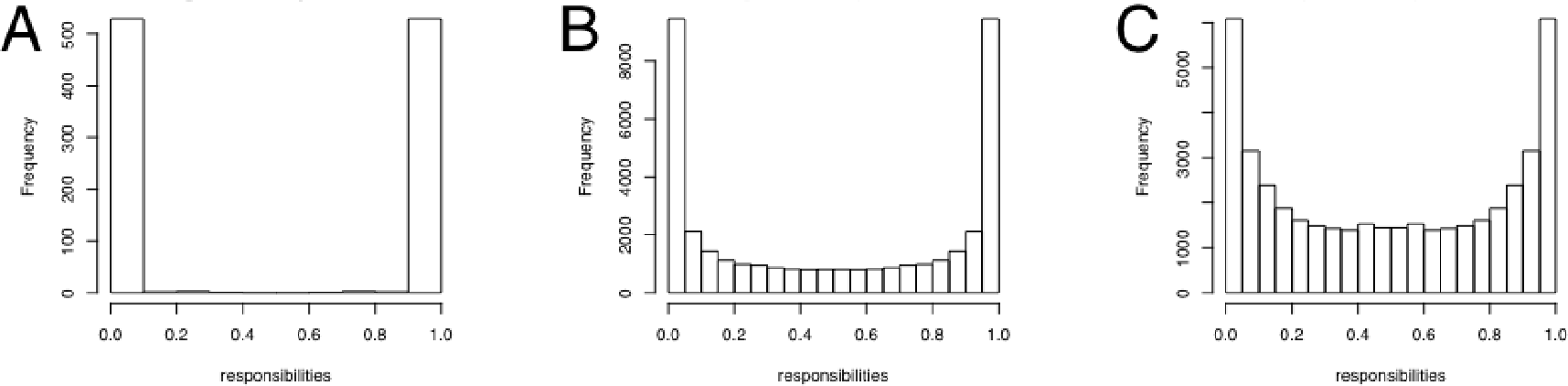
Histograms of responsibilities for Crop-Seq (A), Perturb-Seq cell cycle regulators (B) and transcription factors (C).

A more detailed version of the network with E-gene/Cell attachments for the two highest scoring results (*K* = 2 and *K* = 3) are shown in the supplement, Fig. S3-S4.

### 4.2 Combining CRISPR-based perturbation and RNA-seq (Perturb-Seq)

The data sets of Dixit *et al.* (2016) consists of RNA-seq transcriptome read-outs for single cells. We downloaded them from the BROAD single-cell portal (https://portals.broadinstitute.org/single_cell) and used the log transformed counts per 10000 normalized expression values.

**Cell Cycle Regulators**. Dixit *et al.* (2016) performed knock-out experiments for thirteen cell cycle regulators in K562 cells. After preprocessing the data set consists of 19283 cells and 980 E-genes. Fig. 7 shows the highest scoring M&NEM result (*K* = 2) with mixture weights 46.8% (red) and 53.2% (blue).

**Figure 7:**
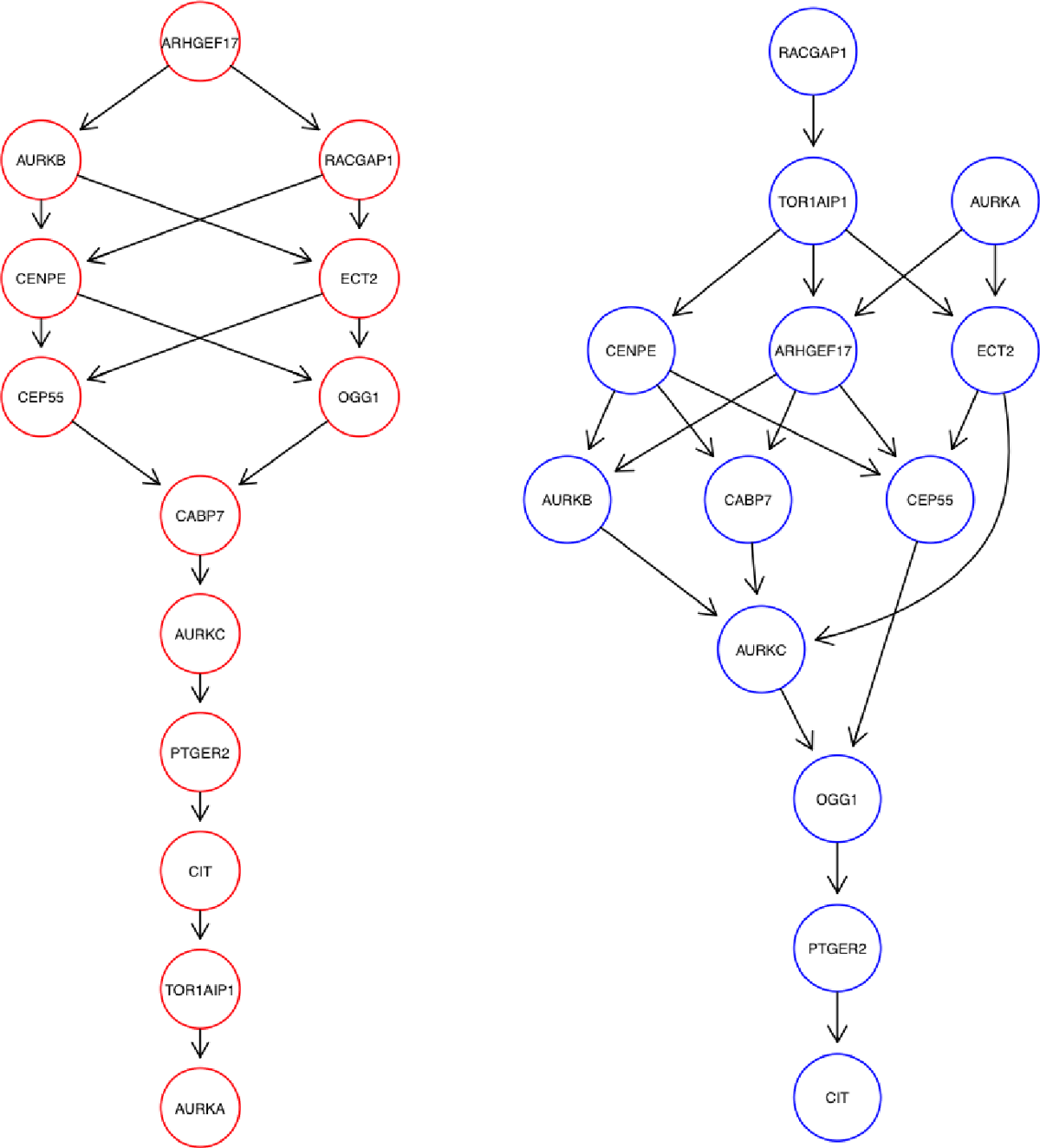
Optimal mixture found for the Perturb-Seq cell cycle regulators (*K* = 2) with mixture weights 46.8% (red) and 53.2% (blue).

Dixit *et al.* (2016) identified the perturbations of PTGER2, CAB7 and CIT as advantageous for proliferation. We found PTGER2 and CIT downstream in both our networks, especially in the heavier blue one, while CAB7 is placed in the middle in both. However, Dixit *et al.* (2016) found a distinct transcriptional phenotype for CAB7, which can explain the different roles in the networks in comparison to PTGER2 and CIT.

Reciprocally, RACGAP1, TOR1AIP1 and AURKA are placed right at the top of the blue network, while their perturbations are identified by Dixit *et al.* (2016) as disadvantageous to proliferation. However, in the other (red) network, only RACGAP1 remains on top and TOR1AIP1 and AURKA are placed right at the bottom. This hints at much more diverse regulatory roles of the latter two and a necessity for RACGAP1 to stay upstream in the network as a key regulator (Imaoka *et al.*, 2015).

Overall the networks differ also in their general shape. While the red network consists of two co-regulating branches, that converge, the blue network is much more inter-connected.

The histogram of responsibilities is shown in Fig. 6, B. The posteriori attachment of cells shows a much softer gradient than for the Crop-Seq data set. While each S-gene in each component has at least one cell which responsibility ≥ 99%, for many cells the responsibilities are between 5% and 95%.

We show a more detailed depictions of the two highest scoring M&NEMs in the supplement, Fig. S5-S6.

**Transcription Factor Interplay**. In a second data set, Dixit *et al.* (2016) performed knock-out experiments for ten transcription factors in K562 cells. The pre-processed data set consists of 22402 cells and 700 E-genes. Fig. 8 shows the optimal network inferred by M&NEM (*K* = 2) with mixture weights of 53.3% (red) and 46.7%.

**Figure 8:**
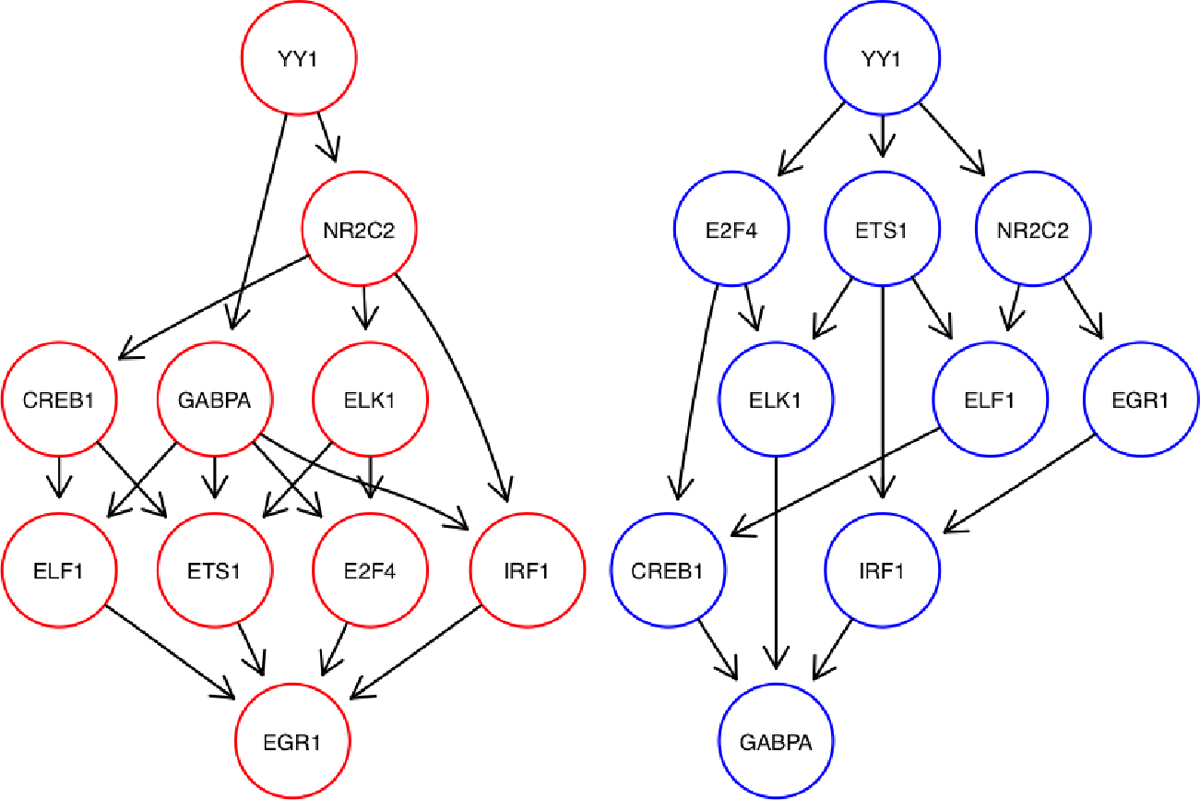
Optimal mixture found for the Perturb-seq transcription factors (*K* = 2) with mixture weights 53.3% (red) and 46.7% (blue).

We identify YY1 as a major regulator for all other genes as it is placed most upstream in both networks. YY1‘s importance as a major transcription factor has been shown before (Tastanova *et al.*, 2016). This is further confirmed as the second highest scoring M&NEM (*K* = 3, supplement, Fig. S8) still places YY1 most upstream in all networks. Similarly, the upstream causal relation of YY1 to NR2C2 is conserved as well.

The other transcription factors mainly switch places in the middle part of the network, except for GABPA and EGR1, who alternately function as the sink node.

Again, the posteriori attachment of cells shows a much softer gradient than for the CROP-seq data set (Fig. 6, A,C). While each S-gene in each component has at least one cell with responsibility ≥ 95%, for many cells the responsibilities are between 20% and 80%.

A more detailed depictions of the two highest scoring M&NEMs is shown in the supplement, Fig. S7-S8.

## 5 Discussion

We have introduced M&NEM, a novel method for the identification of heterogeneous sub populations of single cells with a mixture of networks. M&NEM infers multiple networks from a heterogeneous cell population instead of a single one averaged over the whole population. This additional flexibility allows us to compensate model limitations of the original NEM. M&NEM successfully infers sub populations and the underlaying ground truth mixture of networks in a simulation study under reasonable assumptions.

In our application study, we have investigated three data sets from single cell CRISPR experiments combined with full transcriptomic read-outs. M&NEM confirms known causal interaction and infers novel ambiguous roles for several key regulators (e.g. DOK2), which might be differently regulated in a sub population of cells. We also identify key players like RACGAP1 and YY1, which seem to be necessary for upstream regulation.

Without the use of our model selection to enforce sparseness, our model might lead to over fitting. However, this over fitting might not always be due to noise or technical artifacts, but due to hidden players not perturbed in the data as proposed by Sadeh *et al.* (2013). For example, if we look at the second highest scoring M&NEM for the cell cycle regulators (*K* = 3, supplement, Fig. S6), we see that TOR1AIP1 is placed at the bottom of the blue network with no cells attached and the highest responsibility for a cell at 2%, i.e. almost no information for this placement of the TOR1AIP1 S-gene comes from a cell in which TOR1AIP1 was perturbed. Our hypothesis is, that many E-genes react to AURKA and many E-genes react to CIT, but also many E-genes react to both. Original NEMs cannot model this and it is the exact situation for which Sadeh *et al.* (2013) introduce a hidden player (not perturbed) to account for the diversity of E-genes. In our blue network, TOR1AIP1 is placed to model this scenario and is therefore a stand-in for the unknown hidden player and *not* the actual TOR1AIP1 S-gene (Fig. 9). However, Sadeh *et al.* (2013) use a binomial test based on the binarized data to account for noise, while our model does it in a greedy fashion, which we penalize with our penalized log likelihood ratio. Hence, an integration of the method of Sadeh *et al.*, 2013 into our mixture model to identify hidden players accounting for noise would be an interesting addition.

**Figure 9:**
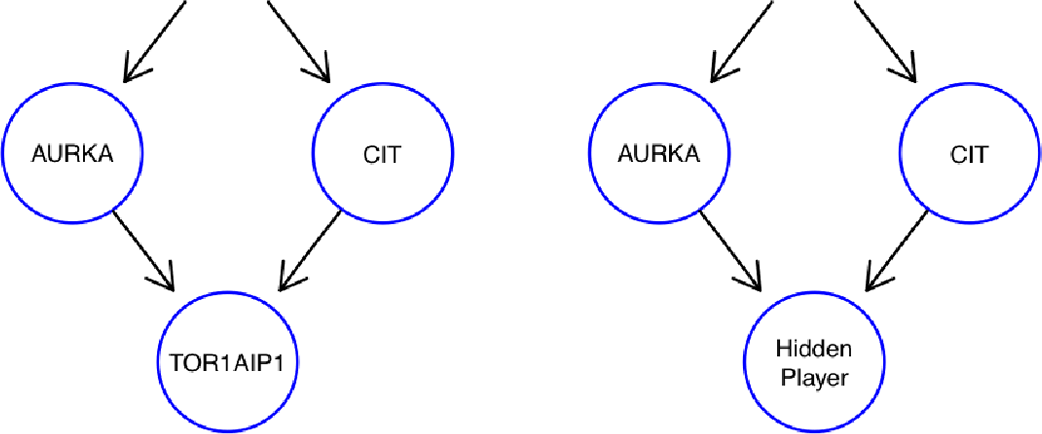
M&NEMs use a degree of freedom to model the possible location of a hidden player not included into the experimental design. The left subgraph shows the M&NEM result with a sink node not supported by any cells and the right subgraph shows our hypothesis for the location of an unperturbed hidden player.

## References

Anchang B., Sadeh M. J., Jacob J., Tresch A., Vlad M. O., Oefner P. J., and Spang R. (2009). Modeling the temporal interplay of molecular signaling and gene expression by using dynamic nested effects models. Proc Natl Acad Sci U S A, 106(16), 6447–6452.

Datlinger P., Rendeiro A., Schmidl C., Krausgruber T., Traxler P., Klughammer J., Schuster L. C., Kuchler A., Alpar D., and Bock C. (2017). Pooled crispr screening with single-cell transcriptome readout. Nature Methods, 14, 297 EP –.

Dempster A. P., Laird N. M., and Rubin D. B. (1977). Maximum likelihood from incomplete data via the em algorithm. Journal of the Royal Statistical Society. Series B (Methodological), 39(1), 1–38.

Dixit A., Parnas O., Li B., Chen J., Fulco C. P., Jerby-Arnon L., Marjanovic N. D., Dionne D., Burks T., Raychowdhury R., Adamson B., Norman T. M., Lander E. S., Weissman J. S., Friedman N., and Regev A. (2016). Perturb-seq: Dissecting molecular circuits with scalable single-cell rna profiling of pooled genetic screens. Cell, 167(7), 1853–1866.e17.

Edgar R., Domrachev M., and Lash A. E. (2002). Gene expression omnibus: Ncbi gene expression and hybridization array data repository. Nucleic Acids Research, 30(1), 207–210.

Friedman N., Linial M., Nachman I., and Pe‘er D. (2000). Using bayesian networks to analyze expression data. J. Comput. Bio., 7, 601–620.

Froehlich H., Fellmann M., Sueltmann H., Poustka A., and Beissbarth T. (2007). Large scale statistical inference of signaling pathways from rnai and microarray data. BMC Bioinformatics, 8, 386.

Froehlich H., Fellmann M., Sueltmann H., Poustka A., and Beissbarth T. (2008). Estimating large-scale signaling networks through nested effect models with intervention effects from microarray data. Bioinformatics, 24(22), 2650–2656.

Froehlich H., Sahin O., Arlt D., Bender C., and Beissbarth T. (2009). Deterministic effects propagation networks for reconstructing protein signaling networks from multiple interventions. BMC Bioinformatics, 10, 322.

Froehlich H., Praveen P., and Tresch A. (2011). Fast and efficient dynamic nested effects models. Bioinformatics, 27(2), 238–244.

Gaudet S. and Miller-Jensen K. (2016). Redefining signaling pathways with an expanding single-cell toolbox. Trends Biotechnol, 34(6), 458–469. 26968612[pmid].

Giancotti F. G. (2014). Deregulation of cell signaling in cancer. FEBS Letters, 588(16), 2558–2570. SI: Tumor suppression.

Imaoka H., Toiyama Y., Saigusa S., Kawamura M., Kawamoto A., Okugawa Y., Hiro J., Tanaka K., Inoue Y., Mohri Y., and Kusunoki M. (2015). Racgap1 expression, increasing tumor malignant potential, as a predictive biomarker for lymph node metastasis and poor prognosis in colorectal cancer. Carcinogenesis, 36(3), 346–354.

Kalisch M. and Bühlmann, P. (2007). Estimating high-dimensional directed acyclic graphs with the pc-algorithm. J. Mach. Learn. Res., 8, 613–636.

MacNeil L. T., Pons C., Arda H. E., Giese G. E., Myers C. L., and Walhout A. J. (2015). Transcription factor activity mapping of a tissue-specific in vivo gene regulatory network. Cell Systems, 1(2), 152–162.

Mao, Hua LeBrun, D. G. Y. J. Z. V. F. L. M. (2012). Deregulated signaling pathways in glioblastoma multiforme: Molecular mechanisms and therapeutic targets. Cancer investigation, 30(1), 48–56.

Margolin A. A., Nemenman I., Basso K., Wiggins C., Stolovitzky G., Dalla Favera R., and Califano A. (2006). Aracne: an algorithm for the reconstruction of gene regulatory networks in a mammalian cellular context. BMC Bioinformatics, 7 Suppl 1, S7.

Markowetz F., Bloch J., and Spang R. (2005). Non-transcriptional pathway features reconstructed from secondary effects of rna interference. Bioinformatics, 21(21), 4026–4032.

Markowetz F., Kostka D., Troyanskaya O. G., and Spang R. (2007). Nested effects models for high-dimensional phenotyping screens. Bioinformatics, 23(13), i305–i312.

Nachman I., Regev A., and Friedman N. (2004). Inferring quantitative models of regulatory networks from expression data. Bioinformatics, 20 Suppl 1, i248–i256.

Pirkl M., Hand E., Kube D., and Spang R. (2016). Analyzing synergistic and non-synergistic interactions in signalling pathways using boolean nested effect models. Bioinformatics, 32(6), 893–900.

Pirkl M., Diekmann M., van der Wees M., Beerenwinkel N., Froehlich H., and Markowetz F. (2017). Inferring modulators of genetic interactions with epistatic nested effects models. PLOS Computational Biology, 13(4), 1–18.

Prasetyanti P. R. and Medema J. P. (2017). Intra-tumor heterogeneity from a cancer stem cell perspective. Molecular Cancer, 16(1), 41.

Sadeh M. J., Moffa G., and Spang R. (2013). Considering unknown unknowns: reconstruction of nonconfoundable causal relations in biological networks. J Comput Biol, 20(11), 920–932.

Siebourg-Polster J., Mudrak D., Emmenlauer M., Raemoe P., Dehio C., Greber U., Froehlich H., and Beerenwinkel N. (2015). Nemix: Single-cell nested effects models for probabilistic pathway stimulation. PLOS Computational Biology, 11(4), 1–21.

Sun, Xiao-xiao Yu, Q. (2015). Intra-tumor heterogeneity of cancer cells and its implications for cancer treatment. Acta Pharmacologica Sinica, 36(10), 1219–27.

Tastanova A., Schulz A., Folcher M., Tolstrup A., Puklowski A., Kaufmann H., and Fussenegger M. (2016). Overexpression of yy1 increases the protein production in mammalian cells. Journal of Biotechnology, 219(Supplement C), 72–85.

Tresch A. and Markowetz F. (2008). Structure learning in nested effects models. Stat Appl Genet Mol Biol, 7(1), Article9.

Wang X., Yuan K., Hellmayr C., Liu W., and Markowetz F. (2014). Reconstructing evolving signalling networks by hidden markov nested effects models. Ann. Appl. Stat., 8(1), 448–480.

